# Tuning and functionalization of logic gates for time resolved programming of bacterial populations

**DOI:** 10.1101/2024.05.14.593743

**Authors:** Leonard E. Bäcker, Kevin Broux, Louise Weckx, Sadhana Khanal, Abram Aertsen

**Affiliations:** Department of Microbial and Molecular Systems, Faculty of Bioscience Engineering, KU Leuven, Kasteelpark Arenberg 23 - bus 2457, 3001 Leuven, Belgium

## Abstract

In order to increase our command over genetically engineered bacterial populations in bioprocessing and therapy, synthetic regulatory circuitry needs to enable the temporal programming of a number of consecutive functional tasks without external interventions. In this context, we have engineered a genetic circuit encoding an autonomous but chemically tunable timer in *Escherichia coli*, based on the concept of a transcription factor cascade mediated by the cytoplasmic dilution of repressors. As proof-of-concept, we used this circuit to impose a time-resolved two-staged synthetic pathway composed of a production-followed-by-lysis program, via a single input. Moreover, via a recombinase step, this synchronous timer was further engineered into an asynchronous timer in which the generational distance of differentiating daughter cells spawning off from a stem-cell like mother cell becomes a predictable driver and proxy for timer dynamics. Using this asynchronous timer circuit, a temporally defined population heterogeneity can be programmed in bacterial populations.

**Graphical abstract:** 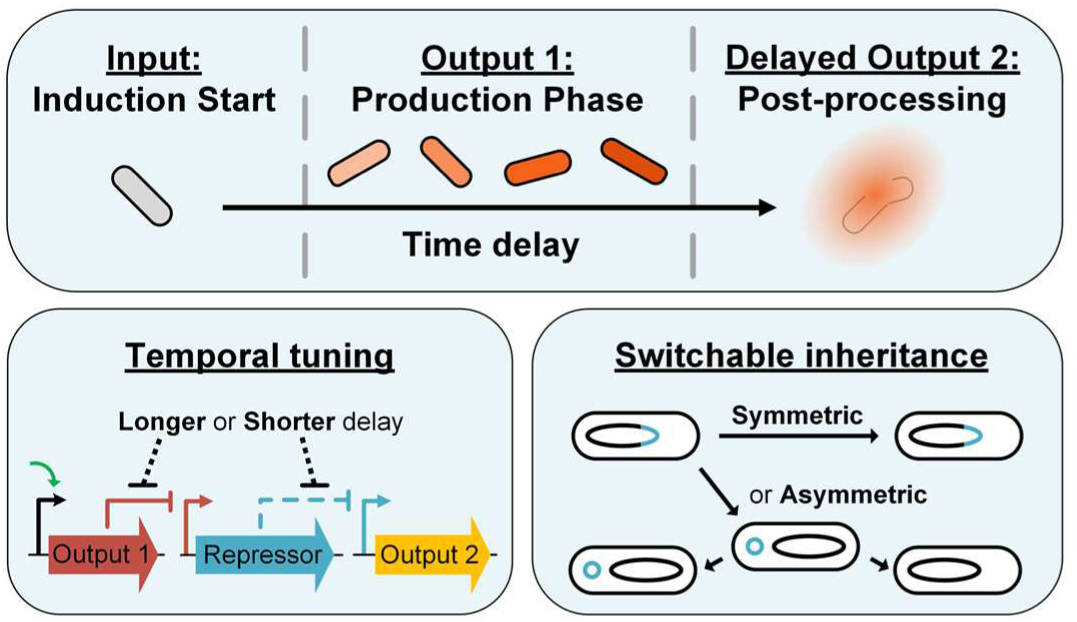

## Introduction

Industrial bioprocessing depends on metabolically engineered microorganisms in bioreactors to (over)produce compounds that they would natively not (or insufficiently) produce (1–3), and is becoming an increasingly important cornerstone in sustainable development and health (4). Indeed, the recent revolution in our capacity to understand and engineer metabolic pathways has resulted in microbial cell factories that can produce a wide variety of biochemical products, including commodities for which no alternative synthesis routes exist (5–8).

However, next to engineering and implementing the metabolic pathways themselves, outstanding challenges for unlocking the full potential of bioprocessing include finding novel ways to increase our overall command of the behaviour of microbial bioreactor populations. As such, synthetic genetic circuitry has already been designed that overlays metabolic pathways with dynamic feedback regulation, whereby the accumulation or depletion of key metabolic intermediates in the cell serves as a trigger to selectively pause up- or downstream branches of the pathway (9,10). Also, quorum sensing dependent circuits have meanwhile been elaborated to properly time and synchronize expression of metabolic pathways throughout bioreactor populations (11,12).

An intriguing class of novel circuitry in this context would be synthetic regulatory systems that aim to autonomously impose a defined timing or succession of gene expression events, as it would consequently allow to spread tasks of bioreactor populations in time. A number of interesting circuits and theoretical models implementing temporal and oscillatory aspects in gene regulatory circuits have already been pioneered (13–18), although usually not designed for the time-resolved programming of gene expression and often accompanied with restrictions on culturing or growth. Nevertheless, a brief succession in gene expression was recently accomplished by Pinto et al. (16), employing cascades of accumulating orthogonal sigma factors, thereby delaying transcriptional responses for up to two doubling times in *Escherichia coli*.

Despite this progress, our means to more elaborately program bioreactor populations and subpopulations with predefined consecutive tasks over several generations is still rather limited. In this study, we therefore devised a timer circuit that is based on cascaded repressors in which the delay is imposed by the gradual dilution of the most downstream repressor. The resulting delay can be chemically tuned and used to autonomously separate a production and subsequent lysis phase in an *E. coli* population. Moreover, this synchronous timer was further engineered to become deterministically asynchronous by imposing recombinase-mediated asymmetric segregation of the DNA fragment encoding the delay-imposing repressor in the cascade.

## Material & Methods

### Strains, media, and cultivation

For all experiments in this study, *Escherichia coli* K12 MG1655 (further referred to as MG1655) and derivatives thereof have been used. All plasmids constructed in this study were maintained in *E. coli* DH10B sAJM.1504 (19). Various strains from the Marionette Sensor Collection, a gift from Christopher Voigt (Addgene Kit #1000000137), were utilized in this study as PCR templates and/or intermediate DNA assembly hosts (19).

For cultivation either Lysogeny Broth (LB; 10 g/L tryptone, 5 g/L yeast extract, 5 g/L NaCl; with 1.5% agar added for agar plates), or AB medium (2 g/L (NH_4_)_2_SO_4_, 7.5 g/L Na_2_HPO_4_ × 2H_2_O, 3 g/L KH_2_PO_4_, 3 g/L NaCl, supplemented with 0.1 mM CaCl_2_, 1 mM MgCl_2_, 0.003 mM FeCl_3_, 10 mg/L Thiamine, 25 mg/L Uracil, 0.2% Casamino acids, and 0.4% glucose) was used. Where needed, the medium was supplemented with the following antibiotics or inducers at the indicated final concentrations: ampicillin (100 µg/mL), chloramphenicol (30 µg/mL), kanamycin (50 µg/mL), tetracycline (10 µg/mL), gentamicin (20 µg/mL), Cuminic Acid (Cuma, 100 µM), Vanillic Acid (Van, ≤ 100 µM), Anhydrotetracycline (aTc, ≤ 200 nM).

All strains were aerobically grown on an orbital shaker at 37°C with the exception for strains harboring the pKD46 (20) helper plasmid or pCP20 (encoding FLP recombinase (21)), which were grown aerobically at 30°C. For stationary-phase cultures, cells were grown for 16-18 h, while for exponential-phase cultures, stationary phase cells were diluted 1/1,000 in fresh medium and grown for an additional 3 to 4 h. When cultures needed to be kept in continuous exponential phase, ca. 2/3 of the culture volume was replaced with fresh pre-warmed medium every hour. In the case of pulsed inductions, after 1 h of growth in the presence of inducer, cells were washed twice with pre-warmed, inducer free medium.

### Plasmid and strain construction

All plasmids created in this study were constructed via Gibson assembly using homology overhangs of 20-40 bp. Gibson assembly master mix was made by 5-fold diluting 5×ISO buffer (0.5 M Tris-HCl pH 7.5, 50 mM MgCl_2_, 1 mM gATP, 1 mM gTTP, 1 mM gGTP, 1 mM gCTP, 50 mM DTT, 5 mM NAD25%, and 25% PEG-8000) into MilliQ and supplementing with 5.3 U/mL T5-exonuclease (NEB), 33.3 U/mL Phusion DNA polymerase (NEB), and 5.3 U/µL Taq DNA ligase (ABclonal). In preparation for the assembly, DNA fragments were first PCR amplified using primers with respective homology overhangs at their 5’end. If needed, DPN1 digestion was performed in Tango buffer, using <1 µL DPN1 restriction enzyme (10 U/µL, Thermo Scientific) and incubating for 1 h at 37°C, followed by heat inactivation (80°C for 20 minutes). The purified DNA fragments were mixed in a ca. 3:1 ratio (insert:vector), and 5 µL of this mix was added to 15 µL aliquots of Gibson assembly master mix thawed on ice. The resulting samples were incubated in a heat block (VWR) pre-heated to 50°C for 1 h, dialysed, and used for electroporation.

For chromosomal replacements and insertions, Lambda Red recombineering was used (20). For the construction of MG1655 AT, the *lac* regulon was first replaced with a *tetA-sacB* cassette (22) in an MG1655 strain harboring the pKD46 plasmid (20). Next, using the *tetA-sacB* counter-selection protocol of Li et al. (22), the *lac::tetA-sacB* region was substituted with a *P_Tet*_-vanR^AM^* construct (based on parts obtained from Meyer et al. (19)). Subsequently, this construct was expanded by transcriptionally fusing an *mCerulean3-frt-Cm^R^-frt* cassette downstream of *vanR^AM^*. Next, the *frt*-*Cm^R^*-*frt* fragment was replaced by a *cymR^AM^-P_CymRC_-T7RNAP-frt-Km^R^-frt* cassette (constructed based on pAJM.657 (19)), after which the region comprising *P_CymRC_-T7RNAP-frt-Km^R^-frt* was replaced by *P_CymRC_-tetR*-*mKate2-P_VanCC_*-*sYFP2-frt-Cm^R^-frt* (based on parts obtained from Meyer et al. (19)), yielding MG1655 AT.

For the construction of MG1655 AAT, *tetR* of MG1655 AT was first replaced with a *bxb1-Gm^R^* cassette flanked with Bxb1 irreversible recombination sites (based on parts obtained from Bonnet et al. (23)). Next, its *P_Tet*_-vanR^AM^*–*mCerulean3-frt-Cm^R^-frt* fragment was replaced by an *frt-Km^R^-frt* cassette, after which this cassette was flipped with pCP20 (21). Finally, the *Gm^R^* marker was replaced by the previously deleted *P_Tet*_-vanR^AM^*– *mCerulean3-frt-Cm^R^-frt* fragment, resulting in MG1655 AAT.

### Lycopene production and quantification

Overnight cultures of MG1655 AT and its derivatives were diluted 1:500 into Erlenmeyer flasks containing 25-100 mL AB medium, and grown at 37°C on orbital shakers for 4 h. Cultures were subsequently normalized to an OD_600_ of 0.2 using pre-warmed AB medium, and 50 mL of culture supplemented with appropriate inducers were further incubated in Erlenmeyer flasks at 37°C. All strains harboring the pTApp plasmid were continuously grown in the presence of 50 μg/mL kanamycin.

For lycopene extraction, 1 mL culture samples were centrifuged for 10 min at 4°C and 3,400 × *g*. The cells were then washed once with 0.085% NaCl and centrifuged again under the same conditions. Next, all supernatant was carefully removed followed by resuspension of the pellet in pure acetone (99.5%). After overnight storage at 4°C, heat extraction was performed for 20 min in a 55°C heat block shaking at 400 rpm, after which adsorption was measurement at 450 nm using a microplate reader (Multiscan FC, Thermo Scientific).

### Microscopy

Time-lapse fluorescence microscopy experiments were performed by placing cells between AB agarose pads (AB medium supplemented with 1.5% agarose) and a cover glass using Gene Frames (Life Technologies), followed by incubation at 37°C using an Okolab cage incubator (Okolab, Ottaviano, Italy).

Cells were imaged using a Ti-Eclipse inverted microscope (Nikon, Champigny-sur-Marne, France), equipped with a 60× Plan Apo λ oil objective, a TI-CT-E motorized condenser and a Nikon DS-Qi2 camera. CFP, YFP and RFP were imaged with triple dichroic (475/540/600 nm) emission filter, and a SpecraX LED illuminator (Lumencor, Beaverton, USA) as a light source.

### Image analysis

Fluorescence microscopy images were analyzed using the DeLTA 2.0 image analysis pipeline (24) for cell segmentation and fluorescence feature extraction, after which the output was further processed using python. In order to accommodate for crowding effects in time-lapse fluorescence experiments, relative fold changes of fluorescence intensities were calculated by dividing the average cellular fluorescence value of an induced condition by the otherwise equally treated basal fluorescence level of the non-induced control. Therefore, values are given as multiples of the non-induced control basal level at the same time point.

### Statistical analysis

GraphPad Prism was used to determine the statistical significance. When appropriate, two-tailed, paired Student’s t-tests and one-way ANOVA with Tukey’s tests were applied. Significance intervals are given as such: *P < 0.05; **P < 0.01; ***P < 0.001; ****P < 0.0001. Best fit values were determined according to the Hill function with uncertainty represented as 95% prediction bands.

## Results and Discussion

### Construction and characterization of an autonomous timer circuit

Since the gradual dilution of repressors has already been shown to impose an intrinsic (and often unwanted) delay in gene expression (25), we coined a synthetic genetic circuit based on a regulatory cascade of three repressors in *Escherichia coli* (Fig. 1A). To avoid promoter leakiness and/or the copy number of the circuit to interfere with the circuits temporal response, we chose (***i***) to employ the high dynamic range P_Tet*_/TetR, P_CymRC_/CymR^AM^ and P_VanCC_/VanR^AM^ promoter/repressor pairs described by Meyer et al. (19) that can also be chemically derepressed (via Anhydrotetracycline or aTc, Cuminic acid or Cuma, and Vanillic acid or Van, respectively), and (***ii***) to chromosomally insert the circuit as a single copy in place of the *E. coli* MG1655 *lac* operon. Finally, in order to make circuit functionalities readily assessable with time-lapse fluorescence microscopy (TLFM), the circuit was equipped with the genes of three spectrally compatible fluorescent protein reporters (i.e. *mKate2*, *mCerulean3*, and *sYFP2*; Fig. 1A). The resulting *E. coli* MG1655 strain harbouring this **<u>a</u>**utonomous **t**imer circuit is further referred to as MG1655 AT.

**Figure 1:**
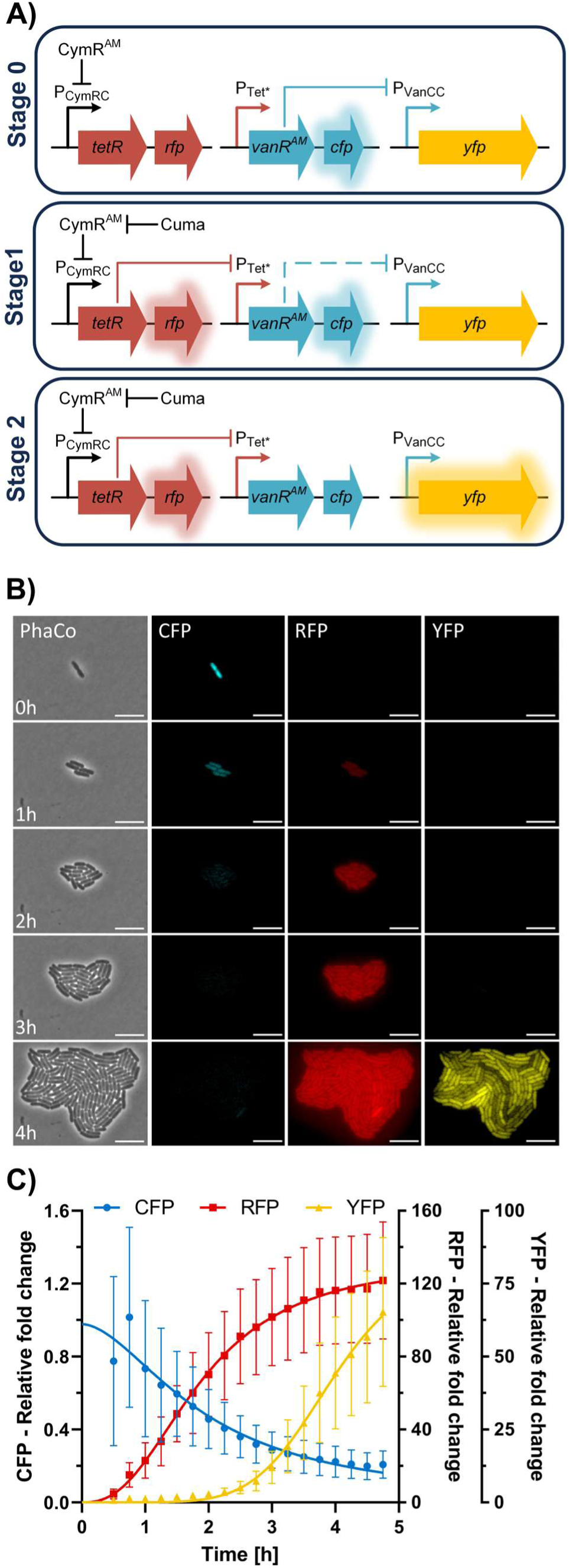
Design and performance of the autonomous timer circuit within *E. coli* MG1655 AT. (A) Schematic illustration of the repressor cascade stages involved in timer progression within MG1655 AT. (B) Representative time-lapse-fluorescence-microscopy image series of MG1655 AT induced with 100 µM Cuma (at t = 0 h) on agarose pads. Scale bars are 10 µm; PhaCo = Phase contrast; CFP = mCerulean3; RFP = mKate2; YFP = sYFP2. (C) Kinetics of the fold change of average cellular CFP, RFP and YFP fluorescence in MG1655 AT cells grown on agarose pads and induced at t = 0 h with 100 µM Cuma relative to similarly grown MG1655 AT cells without induction. Growth of 59 (and 62 for non-induced) single cells into ca. 100-cell microcolonies was monitored. The data depicts means of two independent experiments fitted with a Hill function with error bars representing standard deviation based on cell-to-cell variance.

The transcriptional units of the cascade were arranged such that in the ground state of the circuit (Fig. 1A, Stage 0), the constitutively expressed *cymR^AM^*repressor silences P_CymRC_, which is responsible for the expression of *tetR* and *mKate2* (referred to as *rfp*). Therefore, in the absence of TetR, *vanR^AM^* and *mCerulean3* (referred to as *cfp*) are actively expressed from P_Tet*_. Consequently, in Stage 0, the VanR^AM^ silenced promoter (P_VanCC_) will not express *sYFP2* (referred to as *yfp*). The timer is activated by inducing P_CymRC_ with Cuma (Fig. 1A, Stage 1), resulting in the production of RFP and TetR. The latter represses P_Tet*_, thereby halting *vanR^AM^* expression, and in turn causing the eventual de-repression of P_VanCC_ reported by YFP fluorescence (Fig 1A, Stage 2). Since the initial abundance of the gradually diluting VanR^AM^ determines the actual delay of P_VanCC_ derepression, we employed a translation-maximized *vanR^AM^* ribosome binding site (RBS). Moreover, since leakiness of TetR expression could potentially compromise VanR^AM^ expression levels, *tetR* was fitted with an RBS of moderate strength and an SsrA-tag (expediting degradation of the resulting TetR protein). As such, the cells should only exhibit CFP fluorescence in the non-induced ground state and, upon induction with Cuma, first become red fluorescent, lose their CFP fluorescence, and eventually become yellow fluorescent.

When microscopically assessed on agarose pads (Fig. 1B), MG1655 AT showed no signs of leakiness and presented a delay time of ca. 3 h, as determined by the temporal separation between start of *rfp* expression (Stage 1) and *yfp* expression (progression to Stage 2; Fig. 1B). Likewise, besides the immediate increase of RFP fluorescence, CFP fluorescence started to decline within the first hour of induction (Fig. 1B). Therefore, P_Tet*_ was readily silenced by induction of TetR expression, indicating that the majority of the delay was imposed by the cytoplasmic dilution of VanR^AM^. At the same time, during the first 3 h of timer induction barely any YFP signal was observed, while at ca. 3 h it readily increased.

In order to more precisely assess the temporal dynamics of the individual circuit steps of MG1655 AT, we analysed the time-lapse fluorescence images for average singe cell fluorescence (Fig. 1C). Considering that the maturation time (t_50_) of our RFP (mKate2) is ca. 34 min (26), a significant fluorescence increase within the first 45 min suggests that Stage 1 is initiated readily upon induction. Stage 2 activity is being reported by the extremely fast maturing YFP (SYFP2, t_50_ = 4 min; (26)) and can therefore be estimated with a negligible offset by direct fluorescence readout. At ca. 3 h of induction, the YFP fluorescence starts exceeding the basal level of the non-induced control by 10-fold, while at ca. 4 h the half maximum expression is reached compared to direct Van-induced Stage 2 derepression (which directly derepresses P_VanCC_; cfr. Fig. S1). At the same time, supplementing pads with both Cuma and aTc prevented the (delayed) increase in YFP fluorescence as P_Tet*_-VanR^AM^ expression would not be blocked by the Cuma induced P_CymRC_-TetR (Fig. S1), proving that the delayed YFP increase in Cuma triggered MG1655 AT indeed stems from TetR silenced VanR^AM^ expression and the subsequent cytoplasmic dilution of VanR^AM^. Therefore, the circuit cloned into MG1655 AT reliably imposes time resolved gene expression.

### Functionalizing the autonomous timer circuit

Since our circuit performed as a robust timer, allowing for an autonomous and synchronous delay between two distinct stages, we proceeded with a functional proof-of-concept by imposing an MG1655 AT population with a time-resolved production-followed-by-lysis program via a single input (Fig. 2A). More specifically, a low copy number **t**imer **app**lication plasmid (i.e. pTApp) was constructed encoding (***i***) a P_CymRC_*-crtEBI* circuit, conferring a lycopene production module, that via its P_CymRC_ promoter is coupled to Stage 1 activation, and (***ii***) a P_VanCC_*-M4Lys* circuit, conferring a lysis module, that via its P_VanCC_ promoter is coupled to Stage 2 activation (Fig. 2B). While the *crtEBI* operon stems from *Erwinia herbicola* and enables lycopene production in *E. coli* from its natural farnesyl pyrophosphate intermediate (27), the *M4Lys* (or *ORF_38*) gene stems from the BSPM4 phage and enables lysis independently from the presence of holin or the Sec pathway (28).

**Figure 2:**
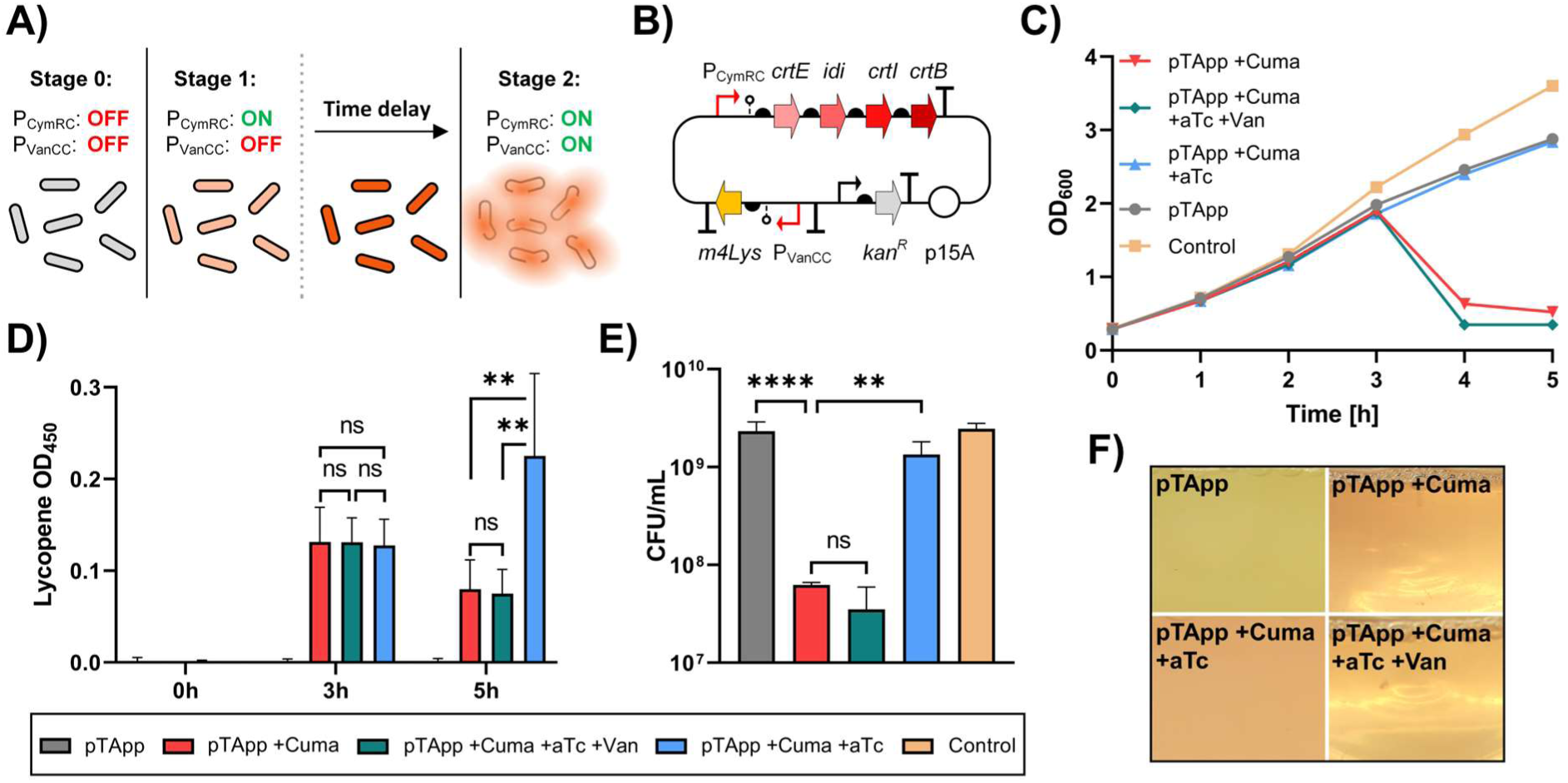
Functionalisation of the repressor cascade-based autonomous timer circuit within *E. coli* MG1655 AT. (A) Schematic illustration of the intended time-resolved production-followed-by-lysis program via a single (Cuma) input. (B) Scheme of the pTApp plasmid harbouring both the P_CymRC_*-crtEBI* circuit (for lycopene production) and the P_VanCC_*-M4Lys* circuit (for subsequent lysis) that serve to functionalize the timer circuit. (C) Kinetics of OD_600_ at indicated time points during batch cultivation of MG1655 AT equipped either without (control) or with pTApp, and the latter being non-induced (pTApp), induced with 100 µM Cuma at t = 0 h (pTApp +Cuma; displaying autonomous timer progression), induced with 100 µM Cuma and 200 nM aTc at t = 0 h (pTApp +Cuma +aTc; fixing the timer at Stage 1), or induced with 100 µM Cuma and 200 nM aTc at t = 0 h and 100 µM Van at t = 3 h (pTApp +Cuma +aTc +Van; imposing transition from Stage 1 to Stage 2 at 3 h). (D) At t = 0 h, 3 h and 5 h of growth, lycopene content of indicated strains was assayed using the plasmid free control as a blank. (E) At t = 5 h viable cell count (CFU/ml) was determined, and (F) representative photographs were taken of indicated strains. For (D) and (E), means and standard deviations of two independent repetitions are shown. For (C) representative results of two independent repetitions are shown.

As intended, when a batch culture of MG1655 AT equipped with pTApp was Cuma induced, it first started to accumulate lycopene for ca. 3 h (Stage 1), after which lysis and reduced viable cell count (Stage 2) autonomously set in (pTApp +Cuma in Figs. 2CDE). In fact, almost complete lysis occurred at 4 h post Cuma triggering, suggesting that time delayed expression of plasmid-based functionalization genes follow the same temporal profile as inferred from the TLFM-assayed fluorescent reporters (Fig. 1), despite the (ca. 10-fold) copy number difference of the pTApp plasmid (∼9 copies/cell; (29)) to the chromosomal YFP reporter. Therefore, the additional P_VanCC_ copies do not significantly titer VanR^AM^ away, and the autonomous timer progression can be used as a temporal queue for pTApp performance. After lysis, the amount of lycopene that could be recovered from cell mass (at t = 5 h; Fig. 2D) was greatly reduced, indicating its release into the culture supernatant.

As a control, MG1655 AT cells were also locked in Stage 1 after Cuma induction by simultaneously providing aTc (pTApp +Cuma +aTc in Figs. 2CDE). This indeed allowed lycopene production to continue (and lycopene levels to increase) past 3 h without being followed by lysis (as P_Tet*_ repression and subsequent P_VanCC_ derepression were abolished). Moreover, when such Stage 1-fixed MG1655 AT cells were forced to proceed to Stage 2 by providing Van after 3 h (pTApp +Cuma +aTc +Van in Figs. 2CDE), they displayed virtually similar lysis and viability dynamics as the autonomously operating pTApp +Cuma population. These same dynamics could also be visually observed as indicated by representative pictures of the cultures after 5 h of incubation in flasks (Fig. 2F). While all Cuma induced cultures had a characteristic pink colour, only the conditions allowing to progress to Stage 2 revealed massive cell lysis.

Since the sequentially expressed functionalisation pathways are located on a small low copy number plasmid harbouring insulated low leakiness promoters form the Marionette Sensor Collection (19,29), the pTApp system is modular and orthogonal to the chromosomal timer. It therefore also allows for gene functions on pTApp to be replaced without the risk of significantly altering timer performance.

### Chemical tuning of the autonomous timer circuit

To further establish the versatility of the autonomous timer circuit, we examined how easily the delay between Stage 1 and Stage 2 could be tuned. In fact, varying the background concentrations of aTc or Van should affect the time it takes to repress P_Tet*_ (directly correlated with aTc concentrations) or to derepress P_VanCC_ (inversely correlated with Van concentrations), respectively (Fig. 3A).

**Figure 3:**
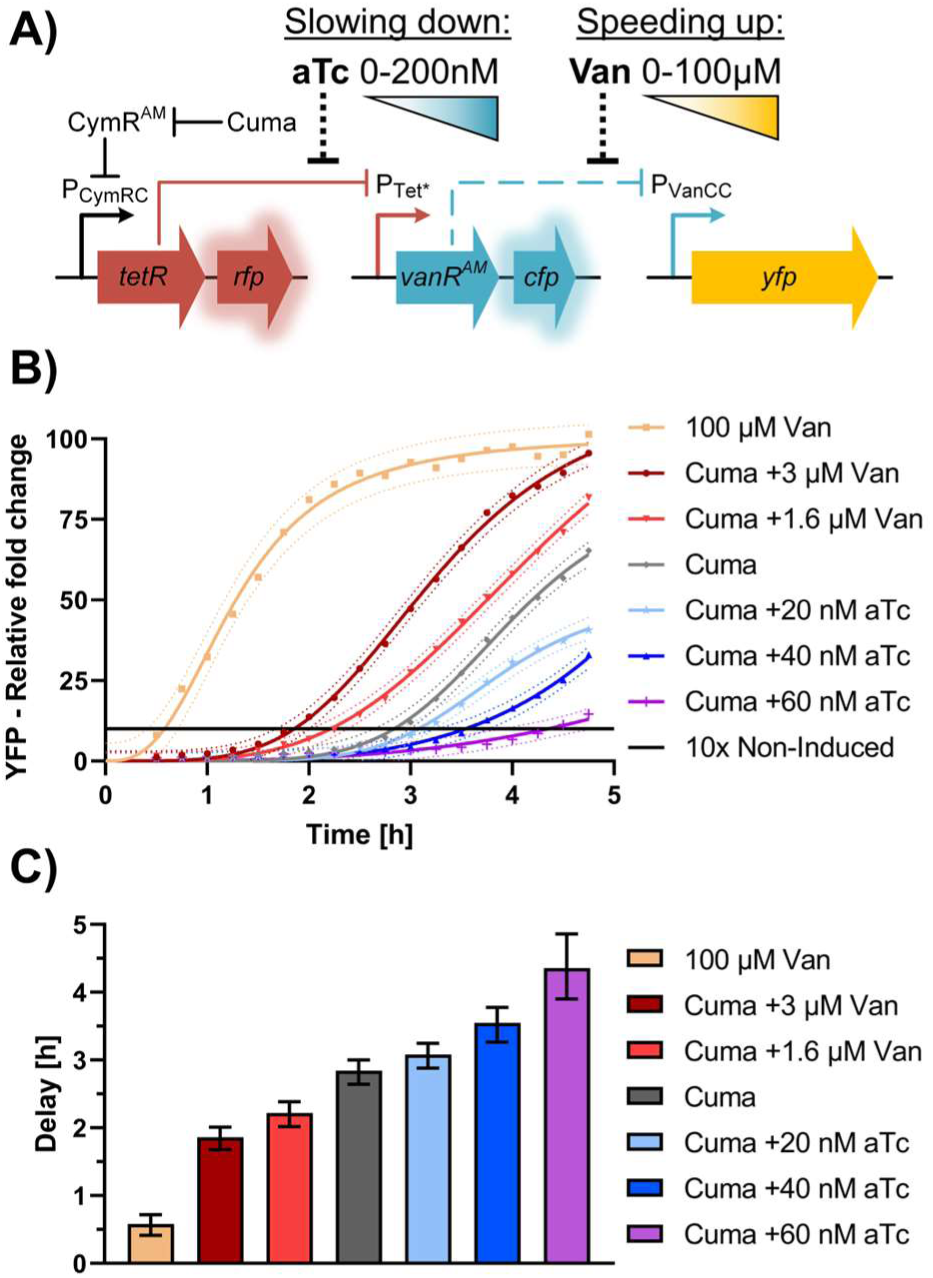
Chemical tuning of the autonomous timer circuit within *E. coli* MG1655 AT. (A) Schematic overview of aTc and Van-based tuning of the autonomous timer circuit. (B) Kinetics of the fold change of average cellular YFP fluorescence of MG1655 AT grown on agarose pads under indicated timer tuning conditions and induced at t = 0 h with 100 µM Cuma, relative to similarly grown MG1655 AT cells without induction. The black line is set at 10-fold the basal YFP intensity level of the non-induced control. The data of two independent repetitions was fitted with a Hill function with errors indicated though 95% prediction intervals. The mean of every datapoint is represented by the respective symbols. For each condition, growth of ca. 60 single cells into ca. 100-cell microcolonies was monitored. (C) Delay in YFP expression estimated by the time it takes for the fitted data in panel B to reach 10-fold the basal YFP intensity level of the non-induced control (i.e. black line in panel B). Error bars indicate the variance by which the upper and lower end of the 95 % prediction interval cross the threshold.

As shown in Fig. 3B, emergence of YFP (as a proxy of Stage 2 activation) can indeed be expedited in the presence of Van and delayed in the presence of aTc. In fact, while 1.6 µM or 3 µM of Van fails to significantly derepress P_VanCC_ when VanR^AM^ is constitutively expressed and thus replenished (cfr. Fig. S1), these concentrations do manage to rapidly attenuate VanR^AM^ activity during the cytoplasmic dilution of this repressor. Overall, the delay (measured as the time after Cuma triggering at which the average cellular YFP expression surpasses the basal level of the non-induced controls 10-fold) of the timer circuit could be tuned between 1.9 h (in the presence of 3 µM Van) and 4.4 h (in the presence of 60 nM aTc; Fig. 3C). Since the doubling time of MG1655 AT remained stable at ca. 38 min throughout all tested conditions, it can also be stated that Stage 2 activation can be chemically tuned to occur anywhere within a range of 3 to 7 cell divisions.

### Construction and validation of an asynchronous autonomous timer circuit

In order to further our abilities to impose a differential but still deterministic timing within bacterial populations, the autonomous timer circuit was adapted so that (***i***) *tetR* became replaced with a likewise SsrA-tagged *bxb1* (encoding the Bxb1 irreversible recombinase with an expedited degradation rate; (23)) and (***ii***) the *P_Tet*_-vanR^AM^-cfp* fragment became flanked with Bxb1 recognition sites (*attB/attP*). The resulting circuit now allows for (pulsed) Cuma-triggering of Bxb1 activity that physically excises the *P_Tet*_-vanR^AM^-cfp* fragment (“*vanR^AM^*-fragment”) from the chromosome and turns it into a circular non-replicating plasmid that inevitably segregates asymmetrically upon cell division (Fig. 4A). Upon each division, cells containing the excised *P_Tet*_-vanR^AM^-cfp* plasmid (referred to as *vanR^AM^*carrier cells) will give rise to a daughter cell lacking the *P_Tet*_-vanR^AM^-cfp* circuit. Since these latter cells thus lack the genetic information to replenish VanR^AM^, they will upon further proliferation experience the progressive dilution of cytoplasmically inherited VanR^AM^ and eventual de-repression of P_VanCC_ (reported by YFP; Fig. 4B). As such, the generational distance of daughter cells from respective *vanR^AM^* carrier cells (i.e. the time since spawning off from a cell expressing VanR^AM^) now is the sole queue for delayed Stage 2 activation (Fig. 4B). The resulting *E. coli* MG1655 strain harbouring this **<u>a</u>**synchronous **<u>a</u>**utonomous **t**imer circuit is further referred to as MG1655 AAT.

**Figure 4:**
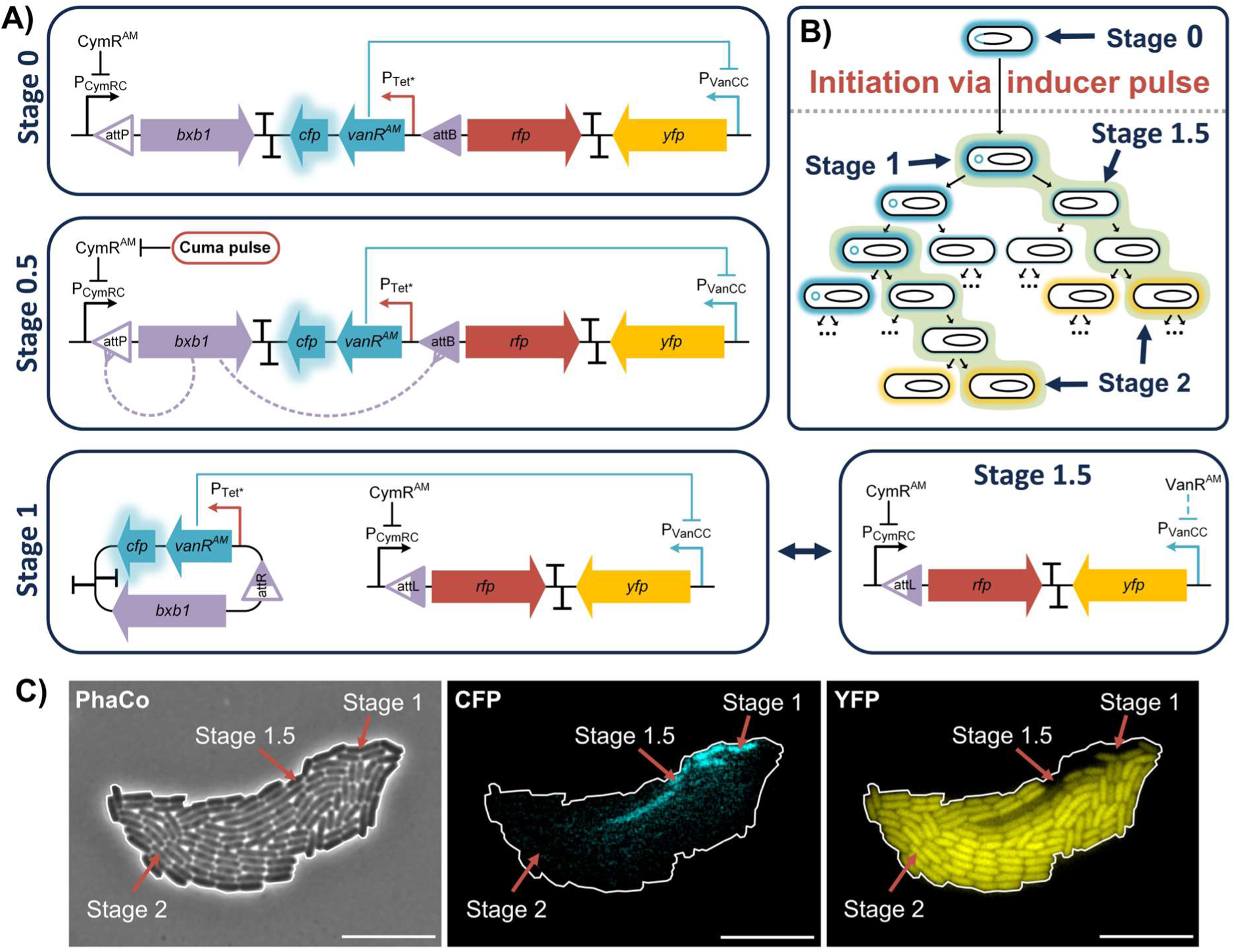
Schematic overview of the asynchronous autonomous timer circuit within *E. coli* MG1655 AAT. (A) Stages involved in the circuit progression. (B) Population based overview of the asynchronous autonomous timer progression in cells spawning off from *vanR^AM^* carrier cells. (C) Microscopy image displaying CFP and YFP fluorescence of a MG1655 AAT microcolony starting from a single cell, pre-induced with 100 µM Cuma for one hour only prior to ca. 5 h of microscopy under non-inducing conditions (see Fig. S2 for full time series). Scale bars are 10 µm; PhaCo = Phase contrast; CFP = mCerulean3; YFP = sYFP2.

When MG1655 AAT cells subsequently became triggered by a short Cuma pulse, they indeed developed into heterogeneous microcolonies that include cells retaining their bright CFP fluorescence over many generations because they asymmetrically inherit the *vanR^AM^*-fragment (underscoring the stability of this episomal fragment), as well as their daughter cells that cytoplasmically dilute their CFP and VanR^AM^ content with every generation and in time become YFP fluorescent (Fig. 4C and Fig. S2). Mapping the CFP/YFP content of cells therefore allows us to infer the following subpopulations: (***i***) *vanR^AM^* carrier cells (Stage 1; displaying high CFP and low YFP), (***ii***) VanR^AM^ diluting cells (Stage 1.5, in between Stage 1 and 2; displaying moderate CFP and low YFP), and (***iii***) VanR^AM^ depleted cells (Stage 2; displaying low CFP and high YFP). Please note that the fraction displaying high CFP and low YFP also contains a number of (Stage 0) cells (ca. 5-10%) that serendipitously failed to Bxb1-excise the *vanR^AM^*-fragment during pulsing, and (passively) remain in the population without partaking in the dynamics. But as shown recently, this fraction could further be suppressed by counter selecting against non-excised cells (30).

Based on this fluorescence gating, a more quantitative analysis of a Cuma-pulsed exponentially growing MG1655 AAT population could be performed (Fig. 5). Since, upon Bxb1-mediated excision, the number of *vanR^AM^* carrier cells remain constant within a population, the VanR^AM^ diluting daughter cells are starting to take over the population. In agreement with the synchronous timer (Fig. 1C), there is a 2 h delay between the first cells experiencing VanR^AM^ dilution reported by decreased CFP fluorescence (as a proxy for entering Stage 1) on the one hand, and subsequently exceeding the basal YFP level of the non-induced control by 10-fold (as a proxy for entering Stage 2) on the other (Fig. 5B).

**Figure 5:**
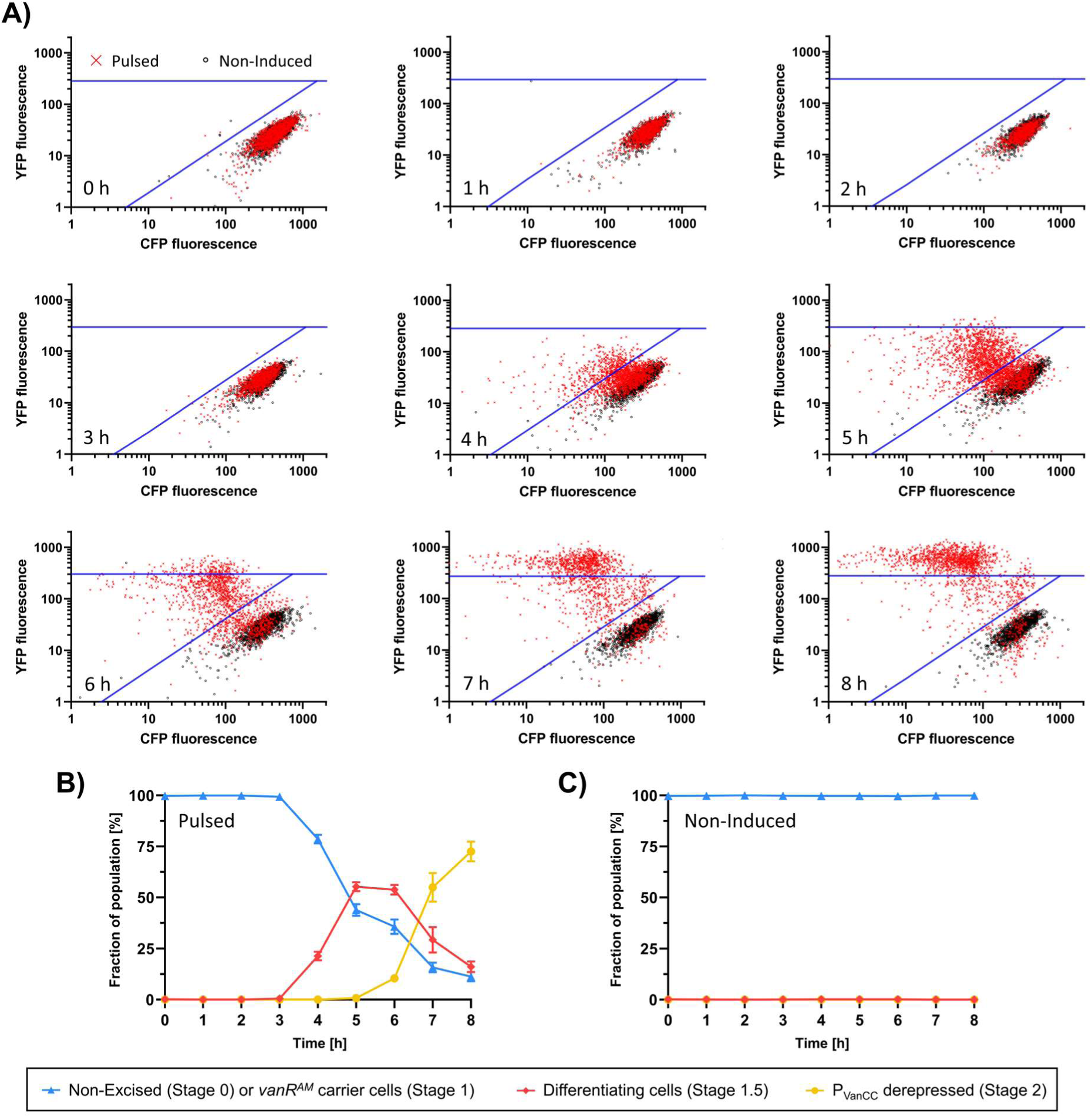
Population dynamics of the asynchronous autonomous timer circuit of *E. coli* MG1655 AAT. (A) Cellular YFP and CFP fluorescence levels of cells sampled at indicated time points from an exponentially maintained culture of MG1655 AAT that was either pulse-induced for the first hour (from 0 h to 1 h) with 100 µM Cuma or non-induced. The data for each condition represents the means and standard deviations of three biological replicates, for which respectively ca. 750 individual cells were analysed per time point. The blue lines indicate the gating used for panels B and C, with the diagonal blue line depicting the 3-fold average YFP to CFP ratio of non-induced cells at the respective time points, and the horizontal blue line depicting the 10-fold average YFP fluorescence of non-induced cells at the respective time points. (B,C) Evolution of the fractions of the gated populations, representing the respective stages of circuit progression, over time for 1 h Cuma (100 µM) pulse-induced (Pulsed, B) or non-induced (C) *E. coli* MG1655 AAT populations.

These results demonstrate that even in a well-mixed environment the pulse-triggered asynchronous timer mediates the creation of temporally defined, and genetically distinct subpopulations over multiple hours. Although in this proof-of-concept fully differentiated cells inevitably take over the population through growth, this aspect could be quenched or alleviated in case Stage 2 would be engineered to impose a more terminal differentiation such as the cell lysis imposed by the pTApp in the previous section or by selecting for the chloramphenicol resistance encoded on the *vanR^AM^* fragment.

## Conclusion

Embedding functional genetic pathways and cargo within intricate regulatory schemes is becoming increasingly important in synthetic biology, as it allows to increase our command over the behaviour of engineered microbial populations. While there are many ways to control successions in gene expression with serial chemical or physical inducers, such inducers are often expensive (31) and/or not always able to properly reach or accumulate at the targeted population when it operates in complex structured environments (such as soil or the human body; (32,33)). In this report, we demonstrate that the intrinsic temporal dynamics of repressor logic gates can be readily engineered and exploited for the autonomous relay of time-resolved instructions to bacterial populations. The elaborated timer-tools allow to synchronously spread tasks over time, and even to asynchronously divide tasks between clonal siblings in a predictable fashion.

In fact, ways to program population heterogeneity in a deterministic and thus predictable manner are scarce. Irreversible recombinases have often been used to invert the orientation of functional genetic elements (17,34–36), or -more recently-to terminate bacterial chromosome replication by excising the origin of replication (37). However, the apparent stability and inherent asymmetric segregation of a functionalized (i.e. providing the VanR^AM^ repressor) Bxb1-excised fragment proves to yield a deterministic motor for cellular differentiation.

In summary, the timer tools designed and characterized in this study provide new handles in programming complex autonomous behaviors in engineered microbial populations that can be applied in bioprocessing, bioremediation and microbial therapy.

## Supplementary data

Supplementary Data are available online.

## Acknowledgments

The authors would like to thank Kristel Bernaerts and Steffen Waldherr for fruitful discussions.

Conceptualization, L.E.B. and A.A.

Methodology, L.E.B. and S.K.

Investigation, L.E.B., K.B. and L.W.

Formal Analysis, L.E.B., K.B. and A.A.

Writing – Original Draft Preparation, L.E.B. and A.A.

Writing – review & editing, L.E.B. and A.A.

Supervision, A.A.

Funding acquisition, A.A.

## Supplementary data

**Figure S1:**
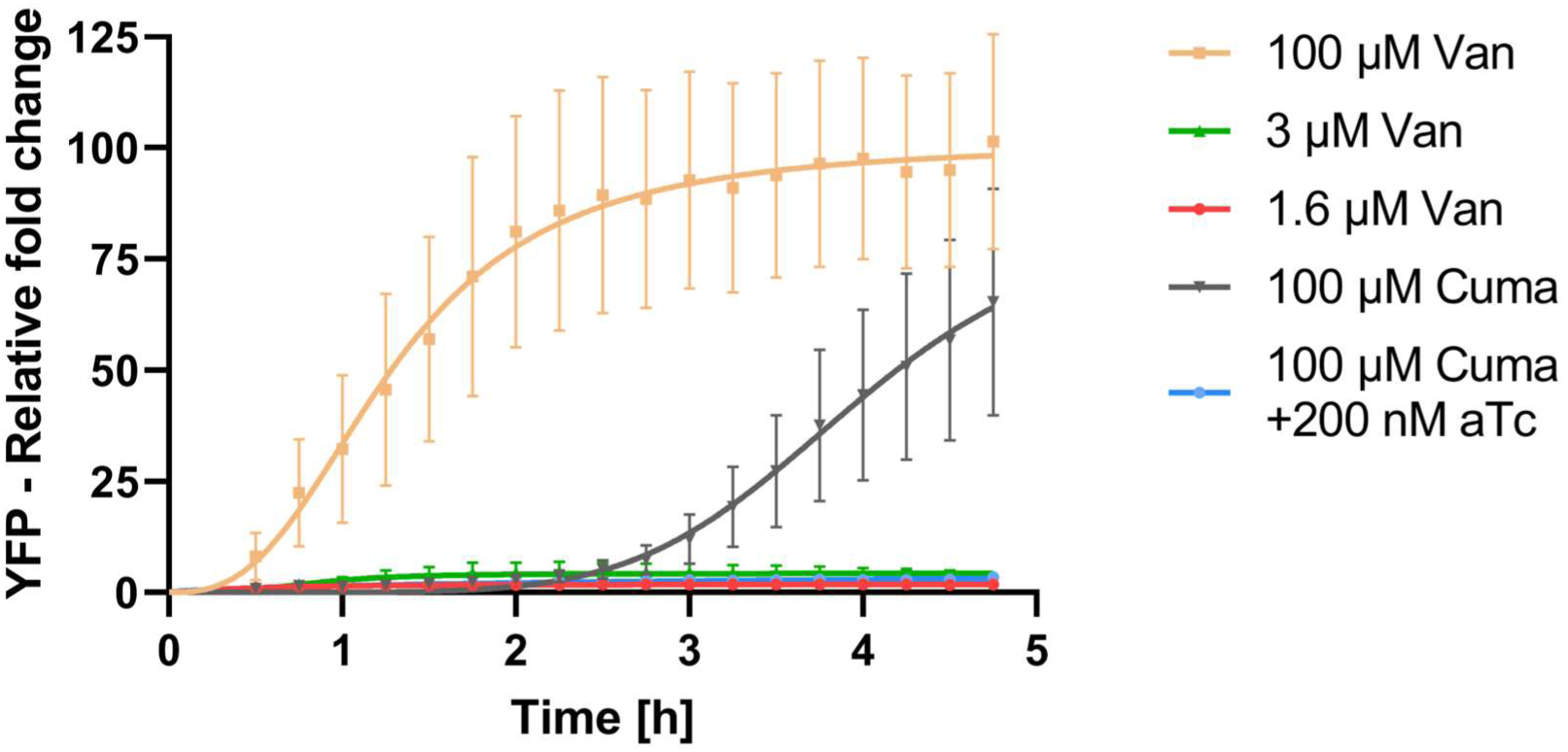
Kinetics of the normalized fold change of average cellular YFP fluorescence in MG1655 AT cells grown on agarose pads and induced at t = 0 h with indicated inducers and concentrations. For each condition, growth of ca. 60 single cells into ca. 100-cell microcolonies was monitored. The data depicts means of two independent experiments fitted with a Hill function with error bars representing standard deviation based on cell-to-cell variance.

**Figure S2:**
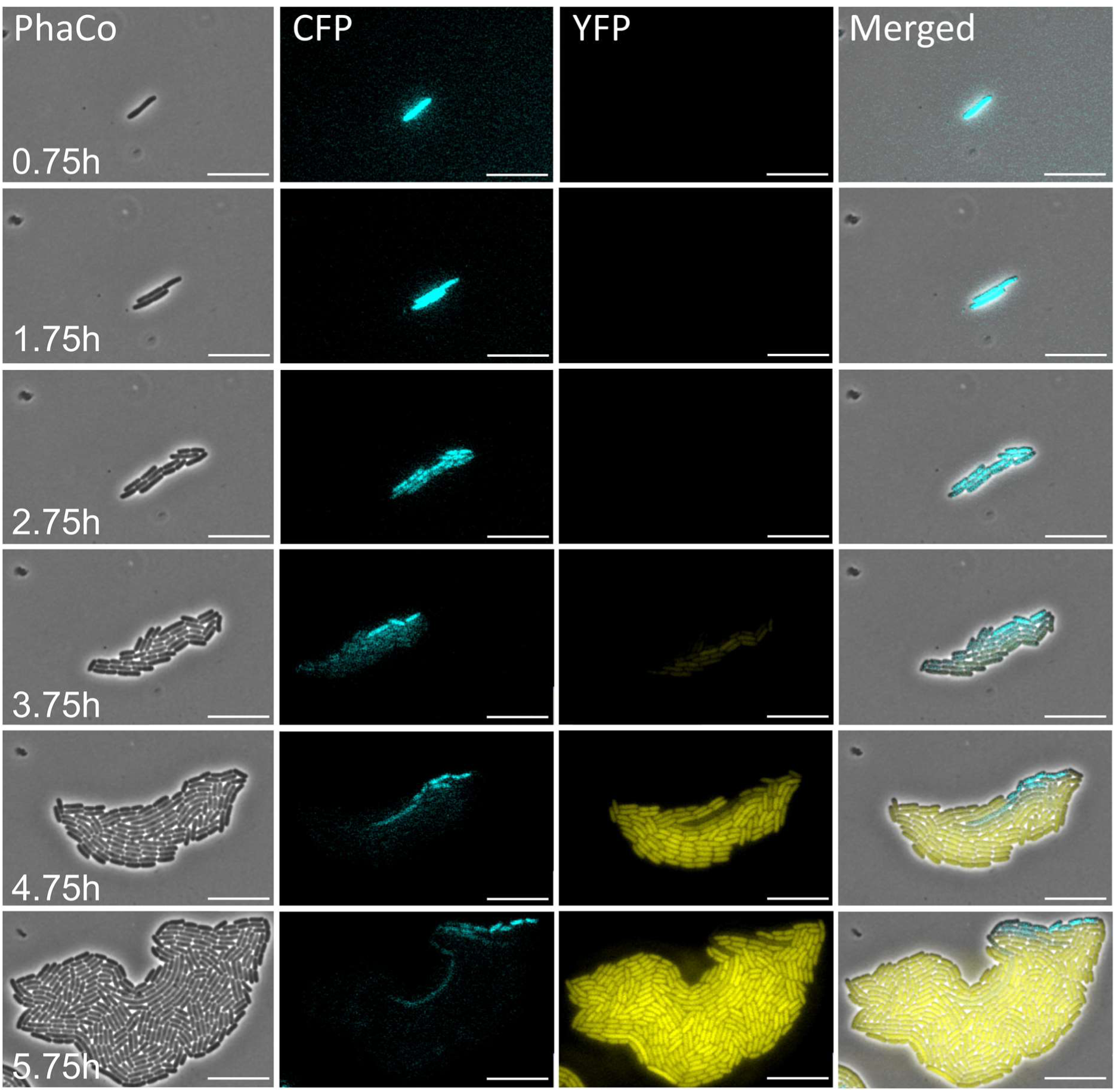
Representative time-lapse-fluorescence-microscopy image series of a MG1655 AAT microcolony starting from a single cell, pre-induced with 100 µM Cuma for one hour only prior to microscopy under non-inducing conditions. Scale bars are 10 µm; PhaCo = Phase contrast; CFP = mCerulean3; YFP = sYFP2

## References

1. Choi, Y.J., Lee, J., Jang, Y.S. and Lee, S.Y. (2014) Metabolic engineering of microorganisms for the production of higher alcohols. Mbio, 5, e01524–01514.

2. Huccetogullari, D., Luo, Z.W. and Lee, S.Y. (2019) Metabolic engineering of microorganisms for production of aromatic compounds. Microb Cell Fact, 18, 41.

3. Majidian, P., Tabatabaei, M., Zeinolabedini, M., Naghshbandi, M.P. and Chisti, Y. (2018) Metabolic engineering of microorganisms for biofuel production. Renew Sust Energ Rev, 82, 3863–3885.

4. Rischer, H., Szilvay, G.R. and Oksman-Caldentey, K.-M. (2020) Cellular agriculture — industrial biotechnology for food and materials. COBIOT, 61, 128–134.

5. Werner, N. and Zibek, S. (2017) Biotechnological production of bio-based long-chain dicarboxylic acids with oleogenious yeasts. WORLD J MICROB BIOT, 33, 194.

6. Yu, X., Liu, F., Zou, Y., Tang, M.-C., Hang, L., Houk, K.N. and Tang, Y. (2016) Biosynthesis of Strained Piperazine Alkaloids: Uncovering the Concise Pathway of Herquline A. J AM CHEM SOC, 138, 13529–13532.

7. Bruckner, S., Haase, R.G. and Schobert, R. (2017) A Synthetic Route to β-Hydroxytyrosine-Derived Tetramic Acids: Total Synthesis of the Fungal Metabolite F-14329. Chem. Eur. J., 23, 5692–5695.

8. Johansson, J. and Curstedt, T. (2019) Synthetic surfactants with SP-B and SP-C analogues to enable worldwide treatment of neonatal respiratory distress syndrome and other lung diseases. J. Intern. Med., 285, 165–186.

9. Liu, D., Mannan, A.A., Han, Y., Oyarzún, D.A. and Zhang, F. (2018) Dynamic metabolic control: towards precision engineering of metabolism. JIMB, 45, 535–543.

10. Lv, Y., Qian, S., Du, G., Chen, J., Zhou, J. and Xu, P. (2019) Coupling feedback genetic circuits with growth phenotype for dynamic population control and intelligent bioproduction. Metab. Eng., 54, 109–116.

11. Gupta, A., Reizman, I.M.B., Reisch, C.R. and Prather, K.L.J. (2017) Dynamic regulation of metabolic flux in engineered bacteria using a pathway-independent quorum-sensing circuit. Nat. Biotechnol., 35, 273–279.

12. Ge, C., Sheng, H., Chen, X., Shen, X., Sun, X., Yan, Y., Wang, J. and Yuan, Q. (2020) Quorum Sensing System Used as a Tool in Metabolic Engineering. Biotechnol. J., 15, 1900360.

13. Weber, W., Stelling, J., Rimann, M., Keller, B., Daoud-El Baba, M., Weber, C.C., Aubel, D. and Fussenegger, M. (2007) A synthetic time-delay circuit in mammalian cells and mice. Proc Natl Acad Sci U S A, 104, 2643–2648.

14. Koseska, A., Zaikin, A., Kurths, J. and García-Ojalvo, J. (2009) Timing Cellular Decision Making Under Noise via Cell–Cell Communication. Plos One, 4, e4872.

15. Toscano-Ochoa, C. and Garcia-Ojalvo, J. (2021) A tunable population timer in multicellular consortia. iScience, 24, 102347.

16. Pinto, D., Vecchione, S., Wu, H., Mauri, M., Mascher, T. and Fritz, G. (2018) Engineering orthogonal synthetic timer circuits based on extracytoplasmic function σ factors. Nucleic Acids Res, 46, 7450–7464.

17. Hsiao, V., Hori, Y., Rothemund, P.W. and Murray, R.M. (2016) A population-based temporal logic gate for timing and recording chemical events. Mol. Syst. Biol., 12, 869.

18. Elowitz, M.B. and Leibler, S. (2000) A synthetic oscillatory network of transcriptional regulators. Nat., 403, 335–338.

19. Meyer, A.J., Segall-Shapiro, T.H., Glassey, E., Zhang, J. and Voigt, C.A. (2019) Escherichia coli “Marionette” strains with 12 highly optimized small-molecule sensors. Nat. Chem. Biol, 15, 196–204.

20. Datsenko, K.A. and Wanner, B.L. (2000) One-step inactivation of chromosomal genes in Escherichia coli K-12 using PCR products. Proc Natl Acad Sci U S A, 97, 6640–6645.

21. Cherepanov, P.P. and Wackernagel, W. (1995) Gene disruption in Escherichia coli: TcR and KmR cassettes with the option of Flp-catalyzed excision of the antibiotic-resistance determinant. Gene, 158, 9–14.

22. Li, X.T., Thomason, L.C., Sawitzke, J.A., Costantino, N. and Court, D.L. (2013) Positive and negative selection using the tetA-sacB cassette: recombineering and P1 transduction in Escherichia coli. Nucleic Acids Res, 41, e204.

23. Bonnet, J., Yin, P., Ortiz, M.E., Subsoontorn, P. and Endy, D. (2013) Amplifying Genetic Logic Gates. Science, 340, 599–603.

24. O’Connor, O.M., Alnahhas, R.N., Lugagne, J.-B. and Dunlop, M.J. (2022) DeLTA 2.0: A deep learning pipeline for quantifying single-cell spatial and temporal dynamics. PLoS Comput. Biol., 18, e1009797.

25. Hooshangi, S., Thiberge, S. and Weiss, R. (2005) Ultrasensitivity and noise propagation in a synthetic transcriptional cascade. Proc Natl Acad Sci U S A, 102, 3581–3586.

26. Balleza, E., Kim, J.M. and Cluzel, P. (2018) Systematic characterization of maturation time of fluorescent proteins in living cells. Nature Methods, 15, 47–51.

27. Das, S., Schapira, M., Tomic-Canic, M., Goyanka, R., Cardozo, T. and Samuels, H.H. (2007) Farnesyl Pyrophosphate Is a Novel Transcriptional Activator for a Subset of Nuclear Hormone Receptors. MOL ENDOCRINOL, 21, 2672–2686.

28. Bai, J., Lee, S. and Ryu, S. (2020) Identification and in vitro Characterization of a Novel Phage Endolysin that Targets Gram-Negative Bacteria. Microorganisms, 8, 447.

29. Shao, B., Rammohan, J., Anderson, D.A., Alperovich, N., Ross, D. and Voigt, C.A. (2021) Single-cell measurement of plasmid copy number and promoter activity. Nat Commun, 12, 1475.

30. Glass, D.S., Bren, A., Vaisbourd, E., Mayo, A. and Alon, U. (2024) A synthetic differentiation circuit in Escherichia coli for suppressing mutant takeover. Cell, 187, 931–944 e912.

31. Ferreira, R.D.G., Azzoni, A.R. and Freitas, S. (2018) Techno-economic analysis of the industrial production of a low-cost enzyme using E. coli: the case of recombinant beta-glucosidase. Biotechnol Biofuels, 11, 81.

32. Jia, X., Li, Y., Xu, T. and Wu, K. (2020) Display of lead-binding proteins on Escherichia coli surface for lead bioremediation. Biotechnology and Bioengineering, 117, 3820–3834.

33. Lim, B., Zimmermann, M., Barry, N.A. and Goodman, A.L. (2017) Engineered Regulatory Systems Modulate Gene Expression of Human Commensals in the Gut. Cell, 169, 547–558.e515.

34. Bowyer, J., Hsiao, V. and Bates, D.G. (2016), 2016 IEEE Biomedical Circuits and Systems Conference (BioCAS), pp. 464–467.

35. Chiu, T.-Y. and Jiang, J.-H.R. (2017) Logic Synthesis of Recombinase-Based Genetic Circuits. Scientific Reports, 7, 12873.

36. Fernandez-Rodriguez, J., Yang, L., Gorochowski, T.E., Gordon, D.B. and Voigt, C.A. (2015) Memory and Combinatorial Logic Based on DNA Inversions: Dynamics and Evolutionary Stability. Acs Synth Biol, 4, 1361–1372.

37. Kasari, M., Kasari, V., Kärmas, M. and Jõers, A. (2022) Decoupling Growth and Production by Removing the Origin of Replication from a Bacterial Chromosome. Acs Synth Biol, 11, 2610–2622.

